# Teaching Python for Data Science: Collaborative development of a modular & interactive curriculum

**DOI:** 10.1101/2021.06.17.448726

**Authors:** Marlena Duda, Kelly L. Sovacool, Negar Farzaneh, Vy Kim Nguyen, Sarah E. Haynes, Hayley Falk, Katherine L. Furman, Logan A. Walker, Rucheng Diao, Morgan Oneka, Audrey C. Drotos, Alana Woloshin, Gabrielle A. Dotson, April Kriebel, Lucy Meng, Stephanie N. Thiede, Zena Lapp, Brooke N. Wolford

**Affiliations:** Department of Computational Medicine & Bioinformatics, University of Michigan; Department of Microbiology & Immunology, University of Michigan; Neuroscience Graduate Program, University of Michigan; Michigan Neuroscience Institute, University of Michigan; Biophysics Graduate Program, University of Michigan; Department of Pathology, University of Michigan; School of Information, University of Michigan; Department of Environmental Health Sciences, University of Michigan; Department of Electrical Engineering & Computer Sciences, University of California, Berkeley; Kresge Hearing Research Institute, Department of Otolaryngology–Head and Neck Surgery, University of Michigan; Michigan Center for Integrative Research in Critical Care, University of Michigan

## Abstract

We are bioinformatics trainees at the University of Michigan who started a local chapter of Girls Who Code to provide a fun and supportive environment for high school women to learn the power of coding. Our goal was to cover basic coding topics and data science concepts through live coding and hands-on practice. However, we could not find a resource that exactly met our needs. Therefore, over the past three years, we have developed a curriculum and instructional format using Jupyter notebooks to effectively teach introductory Python for data science. This method, inspired by The Carpentries organization, uses bite-sized lessons followed by independent practice time to reinforce coding concepts, and culminates in a data science capstone project using real-world data. We believe our open curriculum is a valuable resource to the wider education community and hope that educators will use and improve our lessons, practice problems, and teaching best practices. Anyone can contribute to our educational materials on GitHub.

## Statement of Need

As women bioinformatics trainees at the University of Michigan (U-M), we experience the gender gap in our field first-hand. During the 1974-1975 academic year, women achieved 18.9% of total Bachelor’s degrees in computer and information sciences in the US (National Center for Education Statistics, 2012). By 1983-1984 this peaked at 37.1%, but fell to 17.6% by 2010-2011. We also see this national trend in the training of the next generation of Bioinformaticians at Michigan Medicine. Since accepting its first students in 2001, the U-M Bioinformatics Graduate Program has graduated 66 male and 22 female doctorates as of 2019. This disparity begins at the applicant level; during 2016-2019 the average percentage of females applying directly to the Bioinformatics PhD program was 35.2%, and the average percentage of female applicants listing Bioinformatics as first or second choice in the Program in Biomedical Sciences, U-M’s biomedical PhD umbrella program was 41%.

Previous research on women’s educational experiences in science, technology, engineering, and mathematics (STEM) have produced various explanations for persistent gender disparities (Benbow & Vivyan, 2016). One explanation is that women often experience stereotype threats that negatively influence their math and science performance and deter them from pursuing STEM as a career (Hill et al., 2010). The majority of our organization’s founding graduate students (all women) began coding in our undergraduate careers or later. We wanted to provide a safe place for local high school women to develop confidence in themselves and their computational skills before college, and be exposed to successful women role models in STEM to counter negative stereotypes.

Girls Who Code, a national organization whose mission is to close the gender gap in technology (Saujani, 2015), was founded in 2012. Because of our personal experiences and the paucity of women in our field (Bonham & Stefan, 2017; National Center for Education Statistics, 2012), we began a Girls Who Code student organization at the University of Michigan in 2017. For the past four academic years we have registered annually as a recognized Girls Who Code Club because the national organization provides name recognition, curriculum resources, guidance for a Capstone Impact Project, and a framework for launching a coding club. Participants in the Club attend weekly meetings at the University of Michigan (when the club is run in person rather than virtually), and are thus largely high school women from the Ann Arbor area. In 2019 we launched our own summer program, the Data Science Summer Experience. When held in person, the Summer Experience is hosted in Detroit to provide the opportunity for high school women outside of Ann Arbor to learn coding skills in an inclusive environment.

The national Girls Who Code organization provides a curriculum that teaches website and application development through programming languages like HTML and Java; however, our biomedical science graduate students generally have limited experience with these languages and with web development. In contrast, many of us have extensive experience performing data science using the Python programming language. Data Scientist was rated the #1 job in America by Glassdoor in 2016-2019, #3 in 2020, and #2 in 2021 (Stansell, 2019). Furthermore, Python is the most popular programming language according to the PYPL PopularitY of Programming Language Index (*PYPL PopularitY of Programming Language Index*, n.d.). Therefore, we believe career exploration in data science using the Python programming language will optimally prepare our students for careers that provide financial stability and upward economic mobility. By leveraging the data science expertise of our Club facilitators, we created a specialized curriculum focused on computational data science in the Python programming language.

Girls Who Code encourages participants to learn programming skills while working on an Impact Project website or application throughout the Club (Girls Who Code HQ, 2021). We created an open source Data Science curriculum that teaches the requisite Python and statistics skills to complete a Capstone Project, where students explore, analyze, and present a data set of their choosing. Using this curriculum, we employ participatory live coding as used by The Carpentries, which is an effective method that engages learners (Nederbragt et al., 2020; Wilson, 2016). Using paired activities, our curriculum follows the “I do, we do, you do” didactic paradigm (Fisher & Frey, 2013). We provide open source resources for both in-person and virtual versions of our curriculum, including videos corresponding to each lesson. While we developed this curriculum for our Girls Who Code Club and Summer Experience, we believe that it can be widely used for teaching introductory coding for data science.

## Collaborative Curriculum Development

We assembled a team of volunteers involved in our club to develop a custom curriculum to teach introductory Python for data science via live coding. We chose the content based on what our students would need to learn to complete a small data analysis project and communicate their findings to their peers. We divided the content by topic into Jupyter notebooks for each lesson, with each lesson taking approximately 15-20 minutes to teach via live coding. Every lesson has a corresponding practice notebook with additional exercises on the same content taught in the lesson, but using different data or variables. We used a similar development workflow as the U-M Carpentries curriculum (Lapp et al., 2021). Briefly, we hosted the curriculum notebooks in a public GitHub repository to facilitate collaborative development and peer review using pull requests. In the initial curriculum drafting phase, developers were assigned lesson and practice notebooks to write. Once the draft of a lesson was completed, the writer opened a pull request and asked for review from a different developer. The reviewer then provided feedback and approved the pull request to be merged into the main branch after the writer made any requested changes. This way, more than one person viewed each notebook before it could be incorporated into the public curriculum, which reduced mistakes and ensured higher quality content. While teaching from the curriculum at the first Data Science Summer Experience, instructors took notes on their experience and made revisions afterward. Maintainers continue to monitor the repository and resolve issues as they arise.

Following the onset of the COVID-19 pandemic, we quickly pivoted our club to a virtual format. In preparation for the 2020 Summer Experience, we switched to a flipped classroom style following feedback from our club participants that it was too difficult to follow along live coding via Zoom. We recorded facilitators teaching the lesson notebooks as if they were live coding, then shared them along with a link to the lesson notebooks for students to code along with while watching the videos. Each video shows the Jupyter notebook alongside the facilitator themselves teaching. This format allowed students to learn at their own pace, then ask questions and practice when we met together virtually.

Our Jupyter notebooks and links to the lesson videos can be accessed on GitHub: https://github.com/GWC-DCMB/curriculum-notebooks

## Curriculum

Our curriculum was designed for high school students with no prior coding experience who are interested in learning Python programming for data science. However, this course material would be useful for anyone interested in teaching or learning basic programming for data analysis.

Our curriculum features short lessons to deliver course material in “bite sized” chunks, followed by practices to solidify the learners’ understanding. Pre-recorded videos of lessons enable effective virtual learning and flipped classroom approaches.

### Learning Objectives

Our learning objectives are:

1. Understand fundamental concepts and best practices in coding.
2. Apply data analysis to real world data to answer scientific questions.
3. Create informative summary statistics and data visualizations in Python.

These skills provide a solid foundation for basic data analysis in Python and participation in our program exposes students to the many ways coding and data science can be used to make large impacts across many disciplines.

### Course Content

Our curriculum design consists of 27 lessons broken up into 5 modules that cover Jupyter notebook setup, Python coding fundamentals, use of essential data science packages including pandas and numpy, basic statistical analyses, and plotting using seaborn and matplotlib (Figure 1) (Harris et al., 2020; Hunter, 2007; Waskom, 2021). Each lesson consists of a lesson notebook, used for teaching the concept via live coding, and a practice notebook containing similar exercises for the student to complete on their own following the lesson.

**Figure 1:**
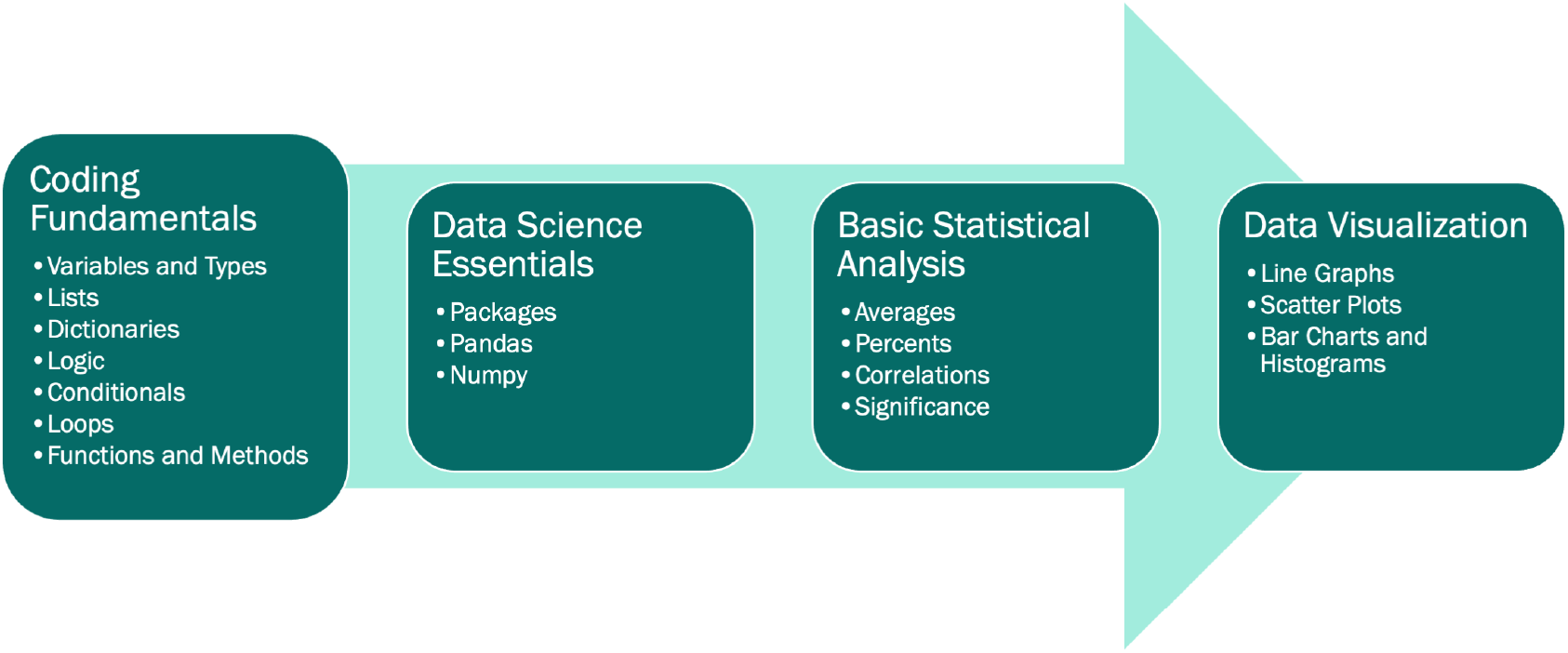
Our lesson modules. All Jupyter notebooks are available on GitHub (https://github.com/GWC-DCMB/curriculum-notebooks).

Each lesson builds on those before it, beginning with relevant content reminders from the previous lessons and ending with a concise summary of the skills presented within. As they progress through the curriculum, the students begin simultaneously working on a data science project using a real world dataset of their choosing. While more time is dedicated to lessons early in the program, the formal curriculum tapers off until the students are solely applying their skills to the data science project.

Through this Capstone Project, learners gain practical experience with each skill as they learn it in the lessons; including importing and cleaning data, data visualization, and basic statistical analyses.

### Instructional Design

We modeled our instructional design in the style of Software Carpentry (Wilson, 2016).

1. Each lesson begins with a recapping of the relevant core skills presented in the previous lessons.
2. All lessons are designed to be taught via 15-minute live-coding sessions, where students type and run code in their own notebooks along with the instructor in real time. As in Software Carpentry, we find this to be a highly effective method of teaching coding, since students must actively engage with the material and deal with errors and bugs as they come up.
3. Each lesson ends with a summary of core skills presented within the material.
4. Each short lesson is also accompanied by a subsequent 10-minute independent practice, providing further opportunity for practical experience implementing the coding skill at hand and testing students’ understanding of the content.

This curriculum was originally developed for in-person instruction, but the onset of the COVID-19 pandemic necessitated restructuring to a virtual format. To better facilitate virtual instruction, we switched to a flipped classroom style based on feedback from our club participants that it was too difficult to follow along with live coding via Zoom. We provide video recordings of instructors going through the Jupyter notebooks as if they were live coding in the classroom. Students then watch the lessons and complete the practice exercises prior to virtual meetings, in which they have the opportunity to ask questions on material they did not understand and go over the practice exercises. This virtual format is especially beneficial because it 1) allows students to learn at their own pace, and 2) enables dissemination of our curriculum to a wider audience interested in learning introductory Python programming for data science.

For both in-person and virtual instruction, once students have completed the Fundamentals module and reach the Data Science Essentials module they begin simultaneous work on their data science projects. Projects are completed in a paired programming style, where partners take turns assuming the “driver” (i.e. the typer) and “navigator” (i.e. the helper) roles (Hannay et al., 2009). Switching off in this way helps both partners assume equal responsibility for the project workload, but more importantly it enables improved knowledge transfer through peer-to-peer learning.

In addition to our coding curriculum, another key component of our programming is hosting women guest speakers from diverse fields across academia and industry. Our guest speakers come to discuss the journey they have taken to their career paths as well as how they utilize programming and data science in their jobs. These varied perspectives are extremely valuable to our learners as they provide several practical examples of programming careers in the real world, and expose them to successful women in STEM.

### Experience of Use

We have used this curriculum to teach the Data Science Summer Experience and Girls Who Code Club in person in 2019 and virtually in 2020-2021. For the in-person instances, we taught the curriculum through participatory live coding, a technique we learned from The Carpentries where the instructor types and explains the code while the learners follow along in real time. For the virtual instances, we used a flipped classroom approach where the learners explore the material individually before meeting together. Learners watch videos of instructors explaining the material through “live” coding and follow along. Learners then complete a practice notebook corresponding to the lesson. Facilitators then spend the meeting time answering questions and reviewing the core concepts in the practice notebook. For both in-person and virtual instances, we had several facilitators present at each session to answer questions and help learners debug. Furthermore, one or two facilitators were assigned to each project group to help learners define data analysis questions, develop and execute a data analysis plan, visualize and communicate their findings, and troubleshoot coding problems. The culmination of the project is a presentation to peers, facilitators, and family members. Through this process learners gain hands-on experience coding, cleaning data, performing statistical analyses, creating informative data visualizations, and communicating their results to others. Projects have ranged from investigating exoplanets to studying the genomics of psoriasis.

We credit the success of our curriculum not only to the skill of the instructors, but also to the way we organized and executed the lessons and project:

1. The instructors and learners used Google Colaboratory (Colab) to write and execute code in Jupyter notebooks. We chose this option because learners do not have to install any programs to use Google Colab, and you can easily open and edit the Jupyter notebooks from GitHub. When meeting in person, most learners use Google Chromebooks which have limited programming capabilities, but easy use of a web browser.
2. Assigning facilitators to groups allowed learners to build a more personal connection with their facilitators, making them feel more comfortable asking questions.
3. Group projects were performed using paired programming to allow learners to collaborate and learn from each other.
4. We used the “sticky note” system from The Carpentries by which learners can ask for help by putting up a colored sticky note (or a Zoom emoji in the case of virtual meetings) (Becker, 2016).
5. We exposed the learners to different aspects of data science by bringing in women guest speakers from academics and industry. This allowed them to better put what they were learning into context, think about how they might use the skills they were learning in potential future careers, and exposed them to successful women in STEM.

### Learner experiences

We surveyed learners anonymously after each Club and Summer Experience and found that most felt that their skills in Python programming, problem solving, critical thinking, and collaboration had improved (see Figure 2). Furthermore, for the 2019 instance of Club we asked that all participants take a pre-test and a post-test comprising Python coding questions. While only five participants recorded their post-test, all of them answered more questions correctly on the post-test than the pre-test (range 1-8 more questions correct out of 11).

**Figure 2:**
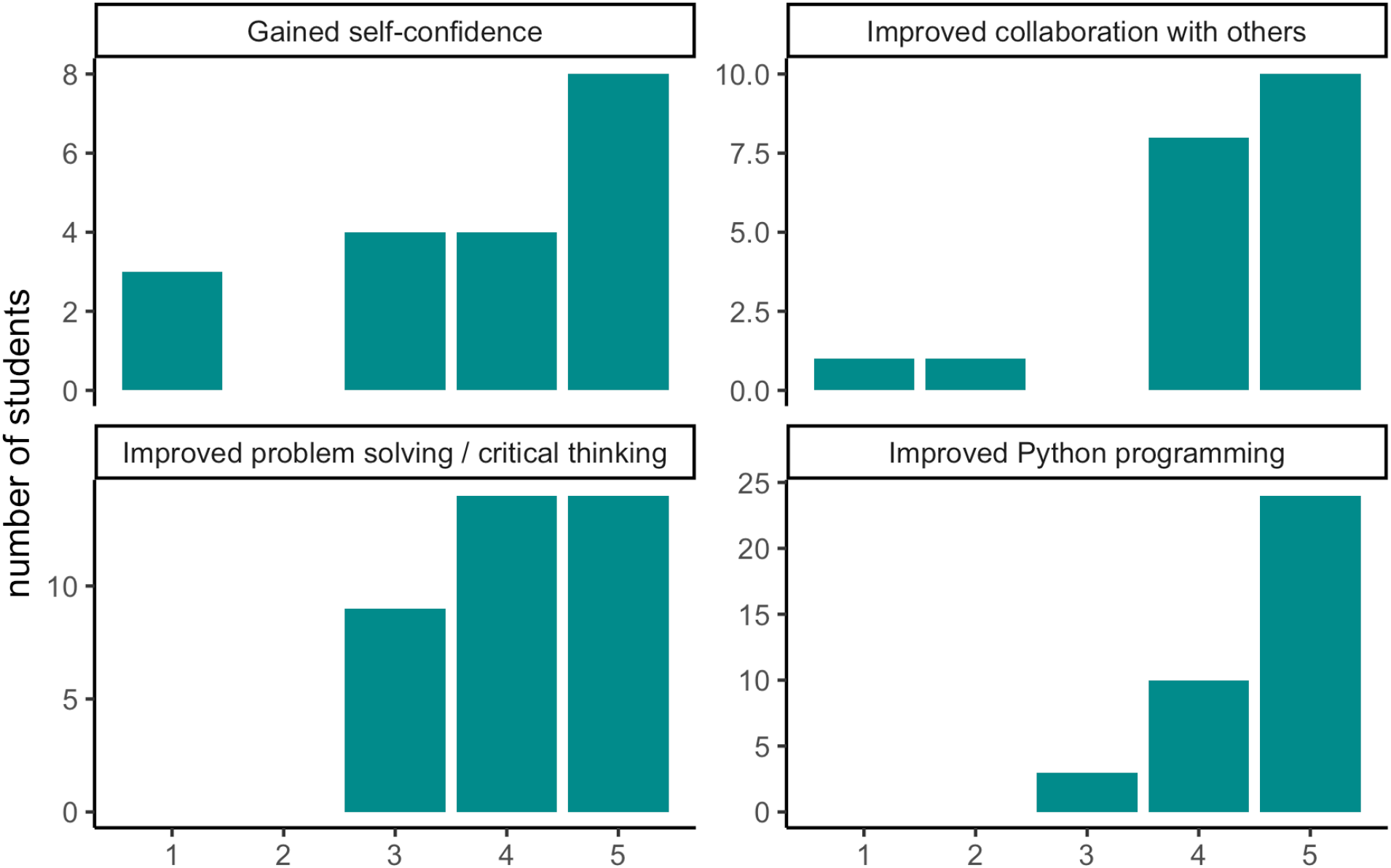
Post-survey responses. Students were asked if they felt that their skills in Python programming, problem solving, critical thinking, and collaboration had improved.

On a 10 question skills assessment, the average increase in correct answers between the first 2019 club meeting and the last 2020 club meeting was 4.2 with a standard deviation of 2.8 (N=5 respondents). Fifty-two percent of the 2019 Detroit Summer Experience in Data Science learners (N=19) did not have experience with Python programming. Afterwards, on a 5 point scale from ‘Not at all’ to ‘Definitely’,’ the average answer for ‘Do you feel like you’ve improved your Python programming skills?’ was 4.6 (standard deviation [s.d.]=1), with 4.2 (s.d.=0.9) for ‘Do you feel like you’ve improved your problem solving and critical thinking skills,’ and 4.0 (s.d.=0.9) for ‘Do you feel like this experience helped you gain self-confidence?’ In a survey of Club and Summer Experience alumni, 75% (N=20) want to pursue a STEM career. 62% (N=21) are still coding. On a 5-point scale from strongly ‘Disagree’ to ‘Strongly Agree,’ the average answer for ‘My participation in GWC impacted my career aspirations’ is 4 (s.d.=0.9), with 4.5 (s.d.=0.6) for ‘Participating in GWC made me feel more confident in analyzing data’ and 3.9 (s.d.=1) for ‘Participating in GWC made me more confident in myself.’ Overwhelmingly, learners’ favorite parts of the program are the guest speakers and the project. These aspects of our curriculum expose them to new fields and allow them to apply their newfound coding skills to asking an interesting question. A 2021 Club learner shared, “I plan to go to college for Computer Science and get a robotics minor when my college offers it. GWC has inspired me to consider pursuing a Masters or PhD in CS as well as take some electives in Data Science.” Five of our 86 alumni have gone on to perform research with U-M faculty members, with one presenting her work at an international conference. In fact, about a third of participants claim that they are now more interested in pursuing a career in computer or data science compared to before their Girls Who Code experience.

## Acknowledgements

We would like to acknowledge our faculty co-sponsors Maureen Sartor & Cristina Mitrea. We appreciate the continued support of U-M DCMB staff and faculty including Julia Eussen, Mary Freer, Linda Peasley, Jane Wiesner, Brian Athey, and Margit Burmeister. We are grateful for the resources provided by the national Girls Who Code organization.

Our programming is made possible by the dedication of past and present Executive Committee members, Club and Summer Experience Facilitators, and Capstone Project mentors including Shweta Ramdas, Alex Weber, Arushi Varshney, Sophie Hoffman, Hojae Lee, Ruma Deb, Saige Rutherford, Michelle McNulty, Bailey Peck, Chloe Whicker, Carolina Rojas Ramirez, Verity Sturm, Zoe Drasner, Sarah Latto, Emily Roberts, Angel Chu, Vivek Rai, Hillary Miller, Ashton Baker, Murchtricia Jones, Lauren Jepsen, Aubrey Annis, Awanti Sambarey, Mengtong Hu, Maribel Okiye, Yingxiao Zhang, and Neslihan Bisgin.

We are grateful for the funding and other support provided to our student organization from the following sponsors: the U-M Department of Computational Medicine and Bioinformatics, the U-M Department of Biostatistics, the U-M Department of Statistics, the U-M Office of Graduate and Postdoctoral Studies, the U-M Endowment in Basic Sciences, the U-M Detroit Center, the U-M Life Sciences Institute, the U-M Office of Research, the Michigan Council of Women in Technology Foundation, DELL Technologies, Cisco Systems, Zingerman’s Delicatessen, the Girls Who Code Support Fund, and anonymous donations from Giving Blue Day 2019.

We also thank the students who have participated in our Club and Summer Experience events.

## Author Contributions

MD, KLS, ZL, and BNW wrote the initial draft of the manuscript. All authors contributed to the curriculum and reviewed the manuscript.

